## Supplementary File 1 for "Blue-shifted ancyromonad channelrhodopsins for multiplex optogenetics"

**Compositions and liquid junction potential (LJP) values of the solutions used in automated patch clamp recording**

|  | **KF** | **NaCl** | **KCl** | **Na aspartate** | **NaNO_3_** | **NMDG Cl** | **CaCl_2_** | **MgCl_2_** | **HEPES** | **EGTA** | **glucose** | **LJP** |
| --- | --- | --- | --- | --- | --- | --- | --- | --- | --- | --- | --- | --- |
| **Internal solution pH 7.2** | 110 | 10 | 10 |  |  |  | 2 | 1 | 10 | 10 |  |  |
| **Standard external solution pH 7.4** |  | 140 | 4 |  |  |  | 2 | 1 | 10 |  | 5 | 7.3 |
| **Standard external solution pH 5.4** |  | 140 | 4 |  |  |  | 2 | 1 | 10 |  | 5 | 7.3 |
| **Na aspartate external solution** |  |  | 4 | 140 |  |  | 2 | 1 | 10 |  | 5 | -4 |
| **NaNO_3_ external solution** |  |  | 4 |  | 140 |  | 2 | 1 | 10 |  | 5 | 6.4 |
| **NMDG Cl external solution** |  |  | 4 |  |  | 140 | 2 | 1 | 10 |  | 5 | 13.5 |
| **KCl external solution** |  |  | 144 |  |  |  | 2 | 1 | 10 |  | 5 | 2.6 |
