## Supplementary File 2 for "Blue-shifted ancyromonad channelrhodopsins for multiplex optogenetics"

**The wavelength positions of the half-maximal amplitude of the long-wavelength slope of the spectrum (λ50) of ChR variants tested in this study**

| **ChR variant** | **λ_50_ (nm)** |
| --- | --- |
| ***Gt*ACR1** | 560 |
| ***Gt*ACR1_S97F** | 574 |
| ***Gt*ACR1_C133M** | 568 |
| ***Gt*ACR1_C237A** | 571 |
| ***Hf*ACR1** | 655 |
| ***Hf*ACR1_F104S** | 629 |
| ***Hf*ACR1_M143C** | 644 |
| ***Hf*ACR1_M143V** | 637 |
| ***Hf*ACR1_A242C** | 646 |
| ***Ans*ACR** | 494 |
| ***Ans*ACR_T90S** | 494 |
| ***Ans*ACR_T90S_D226E** | 506 |
| ***Ans*ACR_M134A** | 495 |
| ***Ans*ACR_M134G** | 496 |
| ***Ans*ACR_M134V** | 502 |
| ***Ans*ACR_D226E** | 506 |
| ***Ans*ACR_A229C** | 496 |
| ***Ans*ACR_A229S** | 496 |
| ***Ft*ACR** | 507 |
| ***Ft*ACR_L96Q** | 489 |
| ***Ft*ACR_M140G_V144A** | 508 |
| ***Ft*ACR_A235C** | 510 |
| ***Ft*ACR_A235S** | 508 |
| ***Nl*CCR** | 482 |
| ***Nl*CCR_F85S** | 487 |
| ***Nl*CCR_S89T** | 485 |
| ***Nl*CCR_M93Q** | 476 |
| ***Nl*CCR_M141A** | 483 |
| ***Nl*CCR_M141C** | 485 |
| ***Nl*CCR_M141G** | 481 |
| ***Nl*CCR_M141V** | 488 |
| ***Nl*CCR_E233D** | 487 |
| ***Nl*CCR_P235I** | 489 |
| ***Nl*CCR_A236C** | 490 |
